# Reinforcement learning discovers new mechanisms of reentry in excitable media

**DOI:** 10.64898/2026.09.08.750287

**Authors:** Thomas M. Bury, Glisant Plasa, José Miguel Romero Sepúlveda, Nicolas Y. Masse, Valentina Romanelli, Leonardo Sacconi, Emilia Entcheva, Gil Bub

## Abstract

The transition from transient excitation to sustained reentry is a fundamental problem in the physics of excitable media. In cardiac tissue, reentry underlies many life-threatening cardiac arrhythmias, yet the pathway to initiation of reentry remains incompletely understood. Here, we formulate reentry initiation as a reinforcement-learning problem in which an agent applies sequences of spatial stimulation patterns while being rewarded for sustained activity and penalized according to the number of stimuli applied. Using cellular automata in one-, two-, and three-dimensional geometries, the agent discovered several mechanisms for generating unidirectional propagation and reentry. These included a previously described mechanism combining superthreshold and subthreshold stimulation, as well as two new mechanisms based entirely on subthreshold stimuli: a sequential mechanism involving stimuli delivered at different locations and times, and a spatial mechanism in which several individually subthreshold sites collectively initiated reentry. In geometries containing boundaries and branches, the learned protocols additionally exploited structural source–sink asymmetries. Optogenetic experiments in cardiac monolayers further demonstrated reproducible induction of unidirectional propagation using the learned spatial subthreshold patterns, while whole-heart experiments provided preliminary evidence that such patterns can shape early propagation in intact tissue. More broadly, we show that reinforcement learning provides a general framework for discovering mechanisms of reentry in arbitrary geometries and generating testable hypotheses about reentry initiation in excitable systems.

## 1 Introduction

Excitable media—including chemical reactions [1, 2], giant honeybee colonies [3], the mammalian cortex [4], forest fires [5], and cardiac tissue [6]—can spontaneously transition between distinct dynamical regimes. Reentry is a self-sustaining rhythm in which an excitation wave repeatedly propagates into and reactivates previously excited regions of the medium. It can take several forms, including circulation around a conduction block [7], rotating spiral or scroll waves [2], and spatiotemporal chaos [8, 9]. Determining how such rhythms are initiated is a fundamental problem in the physics of excitable media because the underlying mechanisms arise from general interactions among excitation, refractoriness, wave propagation, and geometry [9–13]. It is also of particular importance in cardiac tissue, where reentrant rhythms underlie life-threatening arrhythmias [14].

Work on the initiation of reentry dates back to the early twentieth century. Using rings of cardiac tissue, Mines showed that a stimulus delivered near the trailing edge of a previous excitation could produce unidirectional propagation and establish a wave that circulated repeatedly around a closed pathway [15]. This “circus movement” mechanism was further developed by Lewis [16] and later given a mathematical formulation by Wiener and Rosenblueth using a network of connected excitable elements [17]. Selfridge subsequently extended this framework to demonstrate that reentry could also occur as a spiral wave in homogeneous tissue without a fixed obstacle [18]. Since then, a range of initiation mechanisms have been identified, including breakup of a wavefront by external perturbations [2], spatial heterogeneity of refractoriness [19], reduced coupling and structural heterogeneity [12, 13], localized regions of fast conduction [20], structural obstacles [21], and the application of subthreshold pulses [22, 23].

After more than a century of research, predicting which spatial and temporal configurations of localized stimuli are sufficient to initiate reentry remains an open challenge. The problem is especially complex when multiple subthreshold pulses are involved, because propagation depends on their coordinated activity. This scenario is directly relevant to arrhythmias associated with abnormal automaticity, where microscopic cellular or subcellular events combine to generate waves that can evolve into reentrant activity [24–29].

Machine learning methods have become increasingly important in the cardiac sciences, with applications ranging from arrhythmia detection and prediction [30–34] to reconstruction of cardiac excitation dynamics and personalized modeling [35–38]. Most of these approaches are used to analyze data or approximate system dynamics. Reinforcement learning (RL), in contrast, learns sequential decisionmaking strategies through repeated interaction with a dynamical environment. Beyond its successes in robotics, games and control, RL is increasingly being used to automate scientific modelling, reveal biological mechanisms and design sequential interventions in complex dynamical systems where the space of possible strategies is combinatorially large [39–44]. Because the initiation of reentry depends sensitively on the spatial and temporal arrangement of stimuli, and because the number of possible stimulation protocols grows combinatorially with system size, it is well suited for such an approach.

In this article, we formulate reentry initiation as a reinforcement-learning problem and show that RL rediscovers established mechanisms while uncovering new mechanisms across one-, two-, and threedimensional excitable media. We first introduce the cellular-automaton environments and RL framework. We then analyze the mechanisms learned in a onedimensional ring and validate them in continuous reaction–diffusion and cardiac ionic models. Next, we train RL agents in two- and three-dimensional geometries and analyze the distinct stimulation strategies they discover. We show how spatial irregularity of cells can produce orientation-dependent outcomes and how RL can be trained to identify strategies that are robust to changes in orientation and position. Finally, we test one of the spatial subthreshold mechanisms using optogenetic stimulation in cardiac monolayers and intact hearts.

## 2 Reinforcement Learning Framework

Reinforcement learning seeks policies that maximize cumulative reward through repeated interaction with an environment (Fig. 1). At each time step *t*, an agent observes the environment state *s*_*t*_, selects an action *a*_*t*_, and receives a reward *r*_*t*+1_ following the transition to a new state *s*_*t*+1_. In deep RL, the policy is represented by a neural network.

**Fig 1.**
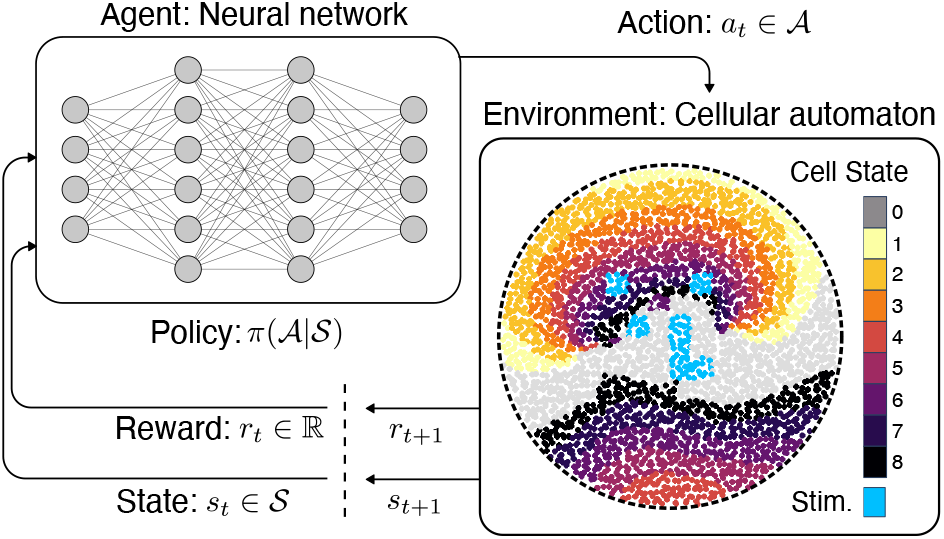
Reinforcement-learning framework for discovering stimulation sequences that induce reentry in excitable media. At time *t*, the agent observes the cellular-automaton state *s*_*t*_ and selects an action *a*_*t*_ according to its policy *π*(*a*_*t* |_ *s*_*t*_). The automaton then advances by one time step, producing the next state *s*_*t*+1_ and reward *r*_*t*+1_. The agent uses the resulting state– action–reward transitions to update its policy. The schematic shows the disk-shaped cellular automaton with a radius of 32 cells, excitation time *E* = 3, refractory time *R* = 5, and an action region located at the center of the domain. Cell color denotes the cellular state: resting (0), active (1–3), or refractory (4–8). Blue cells indicate the stimulus selected by the agent.

In our framework, the environment is an excitable medium modeled by a cellular automaton and the actions are patterns of applied stimuli. The objective is to discover stimulation protocols that initiate reentry while minimizing the number of stimulated cells. This bias toward minimal stimulation discourages trivial strategies in which the agent repeatedly paces the medium to maintain activity rather than initiating reentry. To further promote the discovery of reentry, stimulation is permitted only during an initial time window *t*_stim_. Thereafter, the agent continues to accumulate reward but can no longer intervene. As a result, reentrant rhythms generate large cumulative reward, whereas transient activity rapidly terminates and yields a lower reward.

### 2.1 Cellular Automaton

Since training requires many thousands of simulations, we adopted a computationally efficient cellular automaton model for excitable media [12, 45]. Simulations are carried out on a square lattice. To model spatial irregularity, the position of each cell is perturbed independently along all axes by a random displacement sampled from the uniform distribution *U* [−*ϵ/*2, *ϵ/*2]. The state of cell *j* at time *t* is an integer 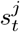 denotes rest; 1, 2, …, *E* are excited states; and *E* + 1, …, *E* + *R* are refractory states. Cells return to the resting state after completing the refractory states. Cells update synchronously. If 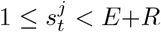, then 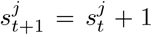; if 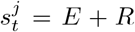, then 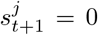. If 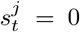, then 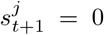 unless the cell is either stimulated by the agent or activated by neighboring cells. A resting cell becomes active if the proportion of neighboring cells within Euclidean distance *ρ* that are in an excited state exceeds the threshold *θ*.

We considered five cellular automaton (CA) geometries spanning one, two, and three spatial dimensions (Fig. 2). Corresponding parameter values are summarized in Table 1. In all geometries, parameters were chosen so that initiation of a traveling wave required stimulation of multiple cells; individual-cell stimuli were subthreshold. For each geometry, we defined an action space consisting of a subset of cells available for stimulation by the agent. The dimensions of the geometries were chosen to be sufficiently large to support sustained reentrant propagation under the corresponding parameter values.

**Table 1.**
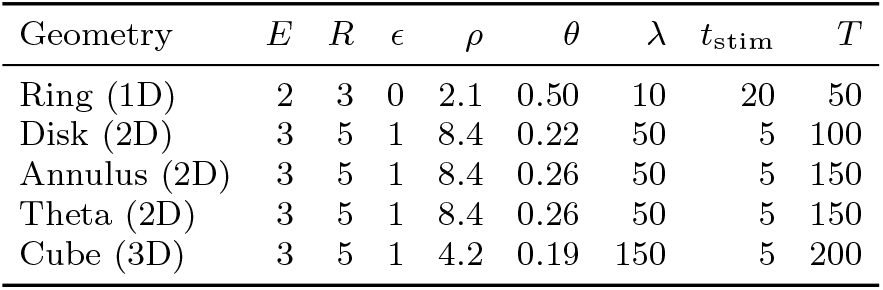
Parameters for each geometry.

**Fig 2.**
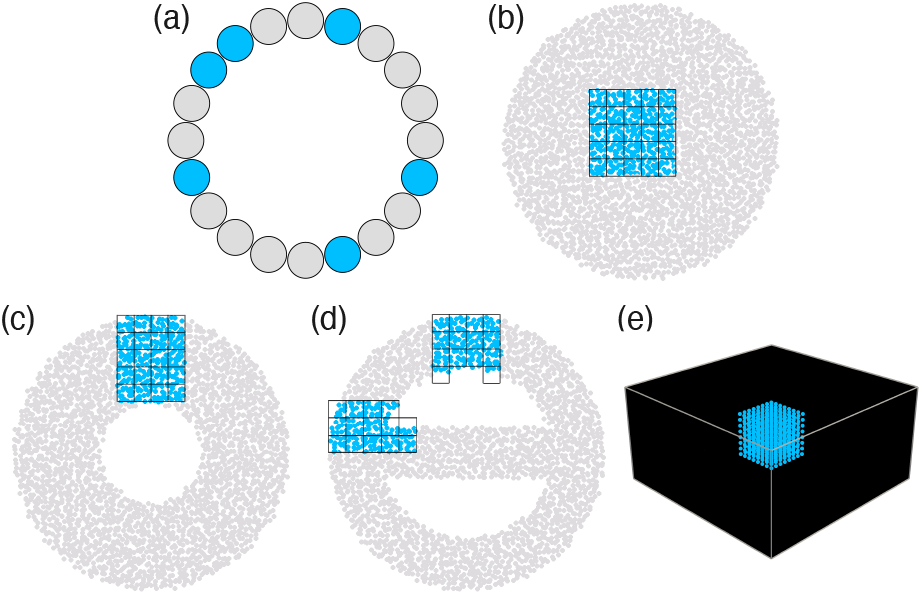
Cellular-automaton geometries used to investigate reentry. (a) One-dimensional ring comprising 20 cells. The agent may stimulate any subset of up to six cells. (b) Twodimensional disk with a radius of 32 cells. Blue indicates the region available for stimulation by the RL agent. This region is partitioned into a 5 *×* 5 grid of stimulus blocks, each comprising 4 *×* 4 cells, and the agent may stimulate any combination of blocks. (c) Two-dimensional annulus with inner and outer radii of 12 and 32 cells, respectively. The stimulation region is partitioned into a 5 *×* 4 grid of 4 *×* 4-cell blocks. (d) Twodimensional theta geometry comprising an annulus connected across its interior by a branch of width 6 cells. The inner and outer radii are 20 and 32 cells, respectively. Two stimulation grids are available: one adjacent to a branch–annulus junction and a second within the annular region. (e) Three-dimensional cube comprising 32 *×* 32 *×* 16 cells. The agent may stimulate any combination of 2 *×* 2 *×* 2-cell blocks within a 4 *×* 4 *×* 4 grid centered in the cube.

To quantify how rarely a single pulse produces reentry, we performed 1,000 simulations per geometry, drawing stimulation patterns uniformly at random from the action space. We then counted the fraction of simulations that resulted in reentry. In the ring geometry, 30.6% of random actions produced reentry; in the disk, 0.1%; in the annulus, 0.0%; in the theta geometry, 2.4%; and in the cube, 0.0%.

### 2.2 Reward Function and Policy Optimization

We set the reward for taking action *a*_*t*_ in state *s*_*t*_ to be

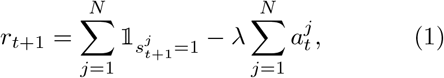

where 1 is the indicator function (equal to 1 if the condition is satisfied, and 0 otherwise), *j* indexes cells, *N* is the number of cells in the geometry, and *λ* is the penalty factor for stimulation. This reward function balances the number of activated cells against the cost of stimulation. For each geometry, we set *λ* such that propagating waves generated by the minimal superthreshold stimulation yield approximately zero net reward. Consequently, the agent can achieve positive return only by discovering stimulation protocols that produce sustained activity more efficiently than repeated direct excitation.

We trained agents using proximal policy optimization (PPO) [46] as implemented in Stable-Baselines3 [47]. We used a learning rate of 3 *×* 10^−4^, batch size of 64, and 10 epochs per policy update. The ring geometry employed a fully connected feature extractor, whereas the two- and three-dimensional geometries used a convolutional feature extractor consisting of three convolutional layers followed by a fully connected layer. PPO used an actor–critic architecture in which the policy and value networks shared the same feature extractor. For evaluation, we report results from the policy that achieved the highest reward during training, which was not necessarily the final policy.

## 3 One-Dimensional Reentry

### 3.1 RL discovery

We trained 10 RL agents on the ring geometry using different random seeds (Fig. 3a). Seven agents learned stimulation protocols that generated sustained reentry; the rest converged to a trivial policy of no stimulation, corresponding to a local optimum of the reward function. The best-performing agent progressively improved during training, evolving from ineffective stimulation patterns to protocols that generated first one then ultimately three concurrent reentrant waves (Fig. 3b,c).

**Fig 3.**
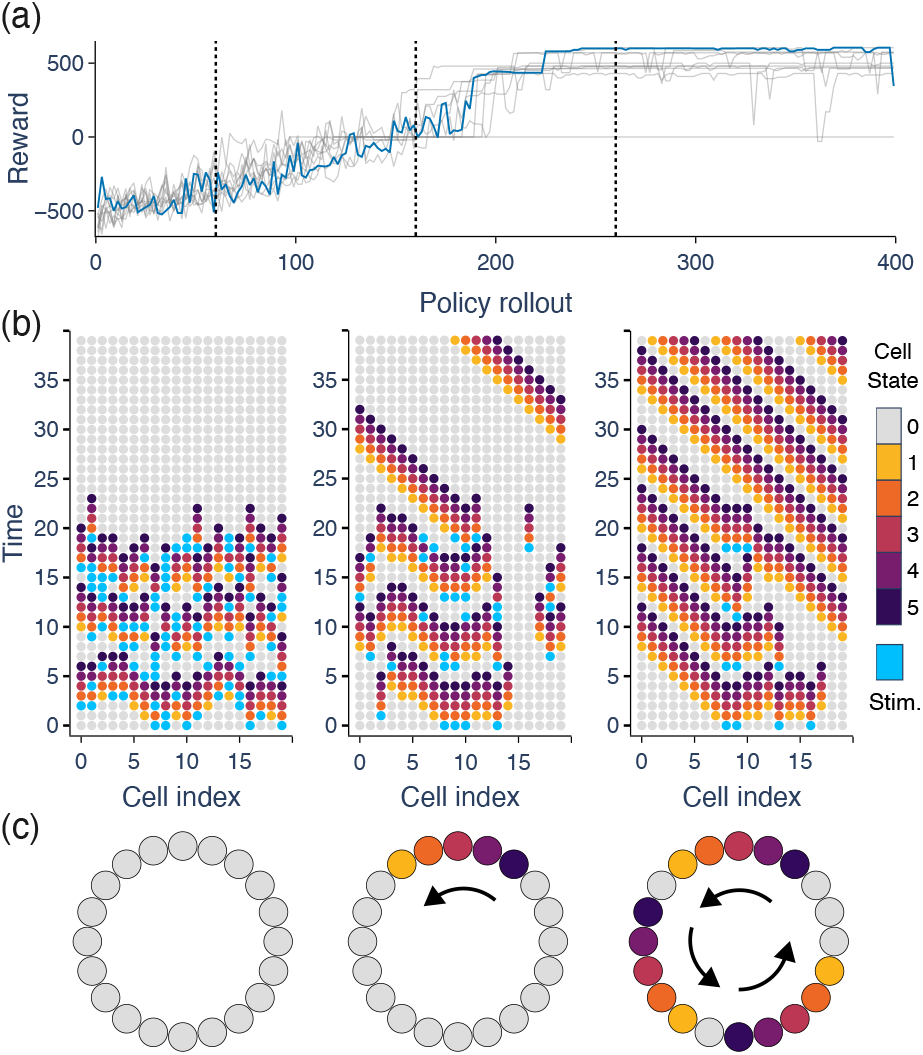
Performance of RL agents during training in the one-dimensional ring environment. (a) Training reward for 10 agents initialized with different random seeds. The bold curve denotes the best-performing agent, and the dashed vertical lines indicate the policy updates illustrated in (b). (b) Representative episodes at policy updates 60, 160, and 260. Cell color denotes the cellular state: resting (0), active (1–2), or refractory (3–5). Blue cells indicate the stimulus selected by the agent. Stimulation is permitted only during the first 20 time steps of each episode. (c) Configurations at the final simulation time containing zero, one, and three reentrant waves, respectively.

Analysis of the learned stimulation protocols revealed two distinct mechanisms for initiating reentry. The first, which we refer to as the *superthreshold– subthreshold* mechanism, combines a superthreshold stimulus with a nearby subthreshold stimulus, creating a localized region of conduction block that converts bidirectional propagation into a unidirectional wave. This mechanism was previously identified by Friedman *et al*. in the FitzHugh–Nagumo model [23]. Its rediscovery by the RL agent validates the framework and demonstrates that known mechanisms of reentry initiation can be learned from reward-driven exploration.

The second, which we refer to as the *sequential subthreshold* mechanism, relies on two subthreshold stimuli delivered at different locations and times. The first stimulus activates a localized group of cells but is insufficient to initiate a propagating wave. When the second stimulus is applied nearby, cells activated by the first stimulus contribute to the combined excitation, allowing the local activity to exceed the threshold for propagation. At the same time, cells at the site of the first stimulus have progressed into refractory states, blocking propagation in that direction. The combination therefore produces a unidirectional traveling wave that subsequently evolves into reentry. To our knowledge, this mechanism has not been previously reported.

### 3.2 Validation in Biophysical Models

To determine whether these mechanisms depend on the discrete nature of the cellular automaton, we tested them in continuous reaction–diffusion models implemented in Myokit [48] (simulation details are provided in the Supplementary Information). The superthreshold–subthreshold mechanism was readily reproduced in the FitzHugh–Nagumo model over a broad range of pulse separations (Fig. 4a,b). The sequential subthreshold mechanism likewise generated unidirectional propagation despite neither stimulus alone being sufficient to initiate a traveling wave (Fig. 4c,d). We further tested both mechanisms in the Fenton–Karma model [49] and the O’Hara– Rudy model [50], where they also produced reentry over a range of pulse separations (Supplementary Figs. 1–2). These results indicate that the mechanisms identified by the RL agent are robust across modeling frameworks ranging from cellular automata to detailed biophysical models.

**Fig 4.**
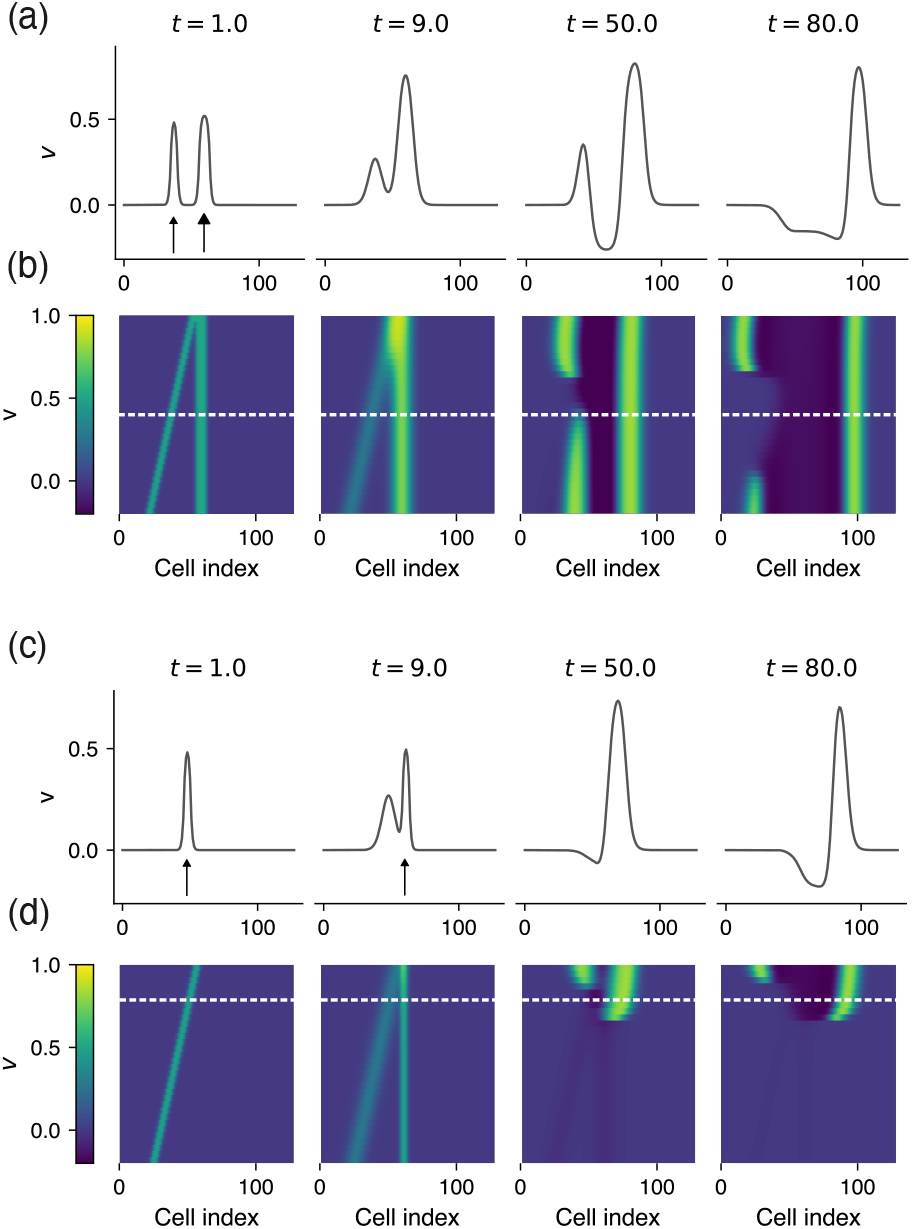
Validation of the superthreshold–subthreshold and sequential subthreshold mechanisms in a 128-cell cable described by the FitzHugh–Nagumo equations. (a,b) Superthreshold–subthreshold mechanism. (a) Voltage profiles at selected times following simultaneous stimulation of two regions separated by 16 cells. One stimulus spans 5 cells and is subthreshold (small arrowhead), whereas the other spans 8 cells and is superthreshold (large arrowhead). Both stimuli are applied for 1 time unit beginning at *t* = 0. (b) Voltage profiles for 32 vertically stacked, independent cables, with the separation between the two stimulus regions ranging from 0 to 31 cells. Each row represents a different stimulus separation, and propagation occurs only in the horizontal direction. The white dashed line marks the cable shown in (a). (c,d) Sequential subthreshold mechanism. (c) Voltage profiles at selected times following two subthreshold stimuli, each spanning 5 cells, applied to regions separated by 8 cells. The second stimulus is delivered 8 time units after the first. (d) Voltage profiles for 32 vertically stacked, independent cables, with the separation between the two stimulus regions ranging from 0 to 31 cells. Each row represents a different stimulus separation, and propagation occurs only in the horizontal direction.

## 4 Two-Dimensional Reentry

### 4.1 Disk geometry

Analysis of the learned stimulation protocols in the disk geometry revealed a third general mechanism, which we refer to as the *spatial subthreshold* mechanism. This mechanism consists of a single stimulus pulse composed entirely of individually subthreshold stimulus sites. Although each stimulus site alone is insufficient to initiate propagation, their asymmetric spatial arrangement generates unidirectional propagation and subsequent spiral-wave formation. This mechanism is closely related to the phenomenon described by Hastings and Sussmann [22], who showed that appropriately combined subthreshold stimuli can initiate spiral waves in the FitzHugh–Nagumo model. Here, the RL agent independently rediscovered this general principle in the cellular automaton model, while identifying distinct spatial arrangements of subthreshold stimulus sites that produce reentry. One solution consisted of four subthreshold stimulus sites arranged in a T-shaped pattern (Fig. 5a). Removing any one site abolished propagation, whereas adding an additional nearby site consistently generated a target wave.

**Fig 5.**
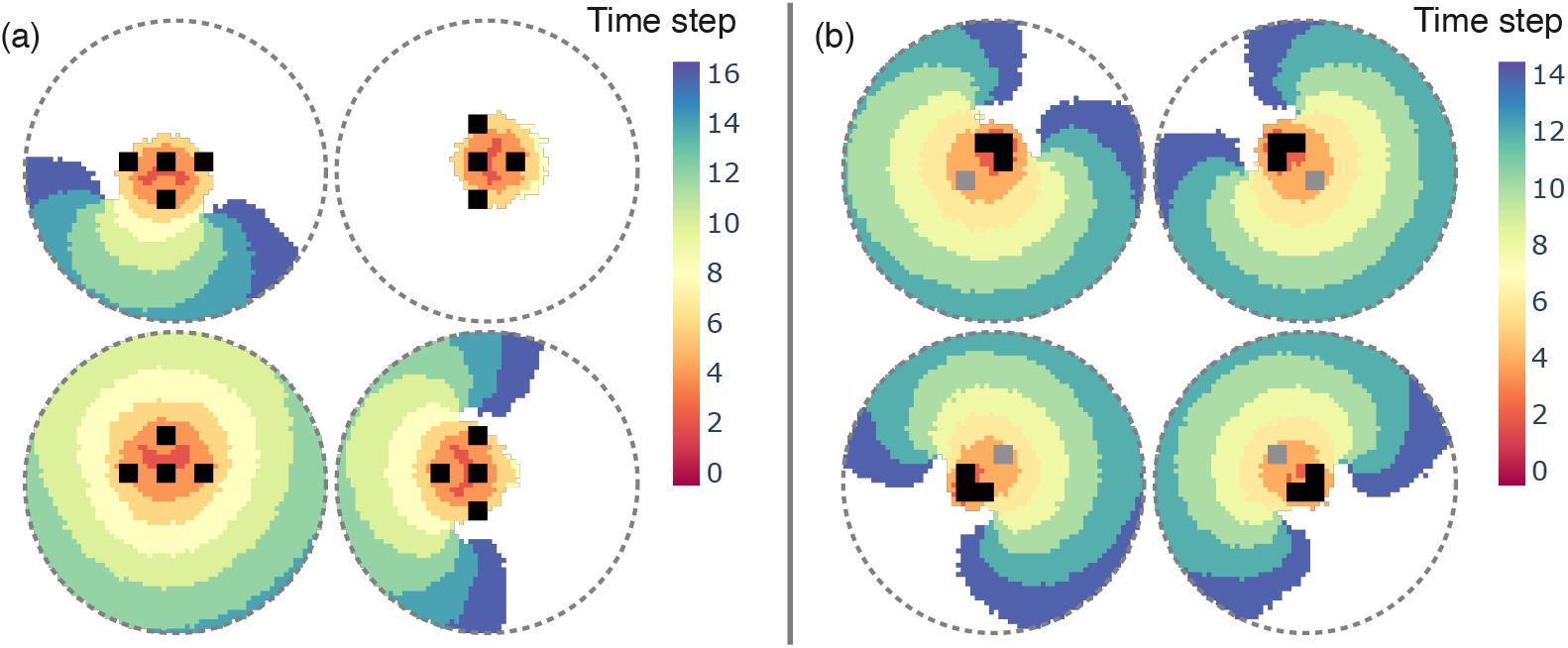
Stimulation patterns learned by the RL agent in the disk cellular-automaton geometry. (a) Activation maps showing wave propagation for four rotations of the spatial subthreshold stimulation pattern (0^°^, 90^°^, 180^°^, and 270^°^). Color indicates the time step at which each cell is first activated; white cells remain inactive, and black cells are stimulated at *t* = 0. Each panel shows the same stimulation pattern applied to the same medium, differing only in its orientation. The resulting dynamics include sustained reentry, a target wave, and propagation block. (b) Activation maps showing wave propagation for four rotations of the sequential subthreshold stimulation pattern in the same disk geometry. Black cells are stimulated at *t* = 0, and gray cells are stimulated at *t* = 2. Sustained reentry occurs for all four orientations.

The spatial subthreshold mechanism is sensitive to the orientation of the stimulus pattern in the cellular automaton model. Rotating the stimulus pattern does not necessarily produce a correspondingly rotated response: depending on its orientation, the same stimulation protocol may produce no propagation, a target wave, or unidirectional propagation (Fig. 5a). This variation in outcome reflects the stochastic spatial arrangement of the cells. We observed the same behavior experimentally (Section 6), motivating the search for stimulation protocols that are robust to changes in orientation and position.

To identify such protocols, we trained the RL agent simultaneously on 16 environments corresponding to four rotations and four spatial translations of the action space. The reward was defined as the average across all environments, encouraging successful initiation of reentry in every case. Under this objective, the agent abandoned the spatial subthreshold mechanism and instead discovered a protocol using the sequential subthreshold mechanism (Fig. 5b). Unlike the spatial subthreshold mechanism, this protocol reliably initiated reentry across all rotations and translations, indicating that the sequential subthreshold mechanism provides a more robust strategy than the spatial subthreshold mechanism in spatially irregular media.

### 4.2 Annulus and Theta Geometries

To investigate reentry initiation in more complex domains, we trained RL agents on annulus and theta geometries, which introduce structural features capable of influencing wave propagation. The theta geometry has previously been used to study distinct varieties of reentrant dynamics arising from the interaction of waves with branching pathways [51]. In the annulus geometry (Fig. 6a,b), agents initiated reentry using both the spatial subthreshold mechanism (a) and the sequential subthreshold mechanism (b). In both cases, however, the learned strategies were tailored to exploit local asymmetries in the geometry. In the examples shown, stimulation occurs near the upper boundary of the annulus. Cells between the stimulation site and this boundary form a relatively small sink and are therefore successfully activated, whereas cells on the opposite side of the stimulus encounter a larger sink, due to the greater number of resting cells between the stimulus and the lower boundary. This asymmetry prevents propagation in one direction while allowing propagation in the other.

**Fig 6.**
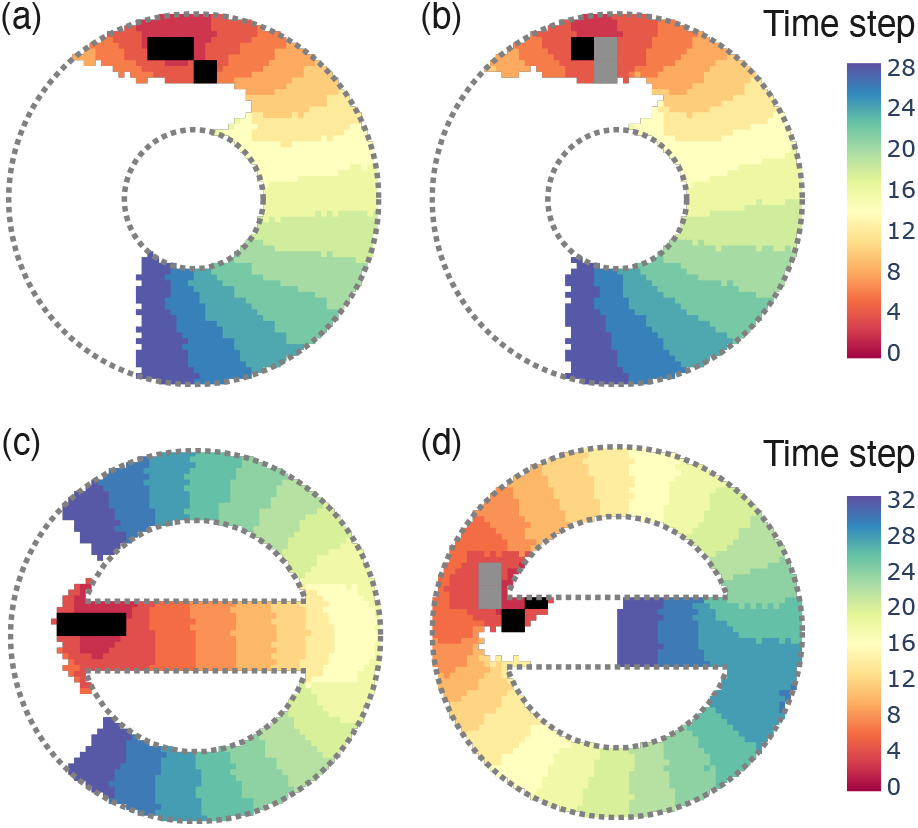
Stimulation patterns learned by the RL agent in the annulus (a,b) and theta (c,d) cellular-automaton geometries. Activation maps indicate the time step at which each cell is first activated. White cells remain inactive, black cells are stimulated at *t* = 0, and gray cells are stimulated at *t* = 1. The learned protocols employ spatial subthreshold stimulation in (a,c) and sequential subthreshold stimulation in (b,d). In each case, the agent exploits the geometry of the domain and the position of the stimulation region relative to nearby boundaries or junctions to generate unidirectional propagation and initiate sustained reentry.

In the theta geometry (Fig. 6c,d), the agent again initiated reentry using both the spatial subthreshold mechanism (c) and the sequential subthreshold mechanism (d), but now exploited the source–sink asymmetry introduced by the branch. For the spatial subthreshold mechanism, stimulation near the branch corner produces preferential activation into the branch, where the available sink is smaller, while propagation into the outer channel is blocked by the larger sink. For the sequential subthreshold mechanism, the direction of propagation is reversed: an initial subthreshold pulse creates refractory block within the branch, while a subsequent pulse boosts activation around the corner where the branch meets the outer channel. Thus, in both geometries, the agent exploits local geometric asymmetries to generate unidirectional propagation and initiate reentry.

## 5 Three-Dimensional Reentry

We next investigated whether stimulation protocols learned in two-dimensional geometries could be extended to three dimensions. Applying the four-site stimulus pattern identified in the disk geometry (Fig. 5a) to the upper surface of the cube, and extending the stimulus to a depth of four cells, initiated a sustained three-dimensional reentrant rhythm (Fig. 7a). Thus, the spatial subthreshold mechanism discovered in two dimensions remains effective in three dimensions, where it gives rise to scroll-wave reentry.

**Fig 7.**
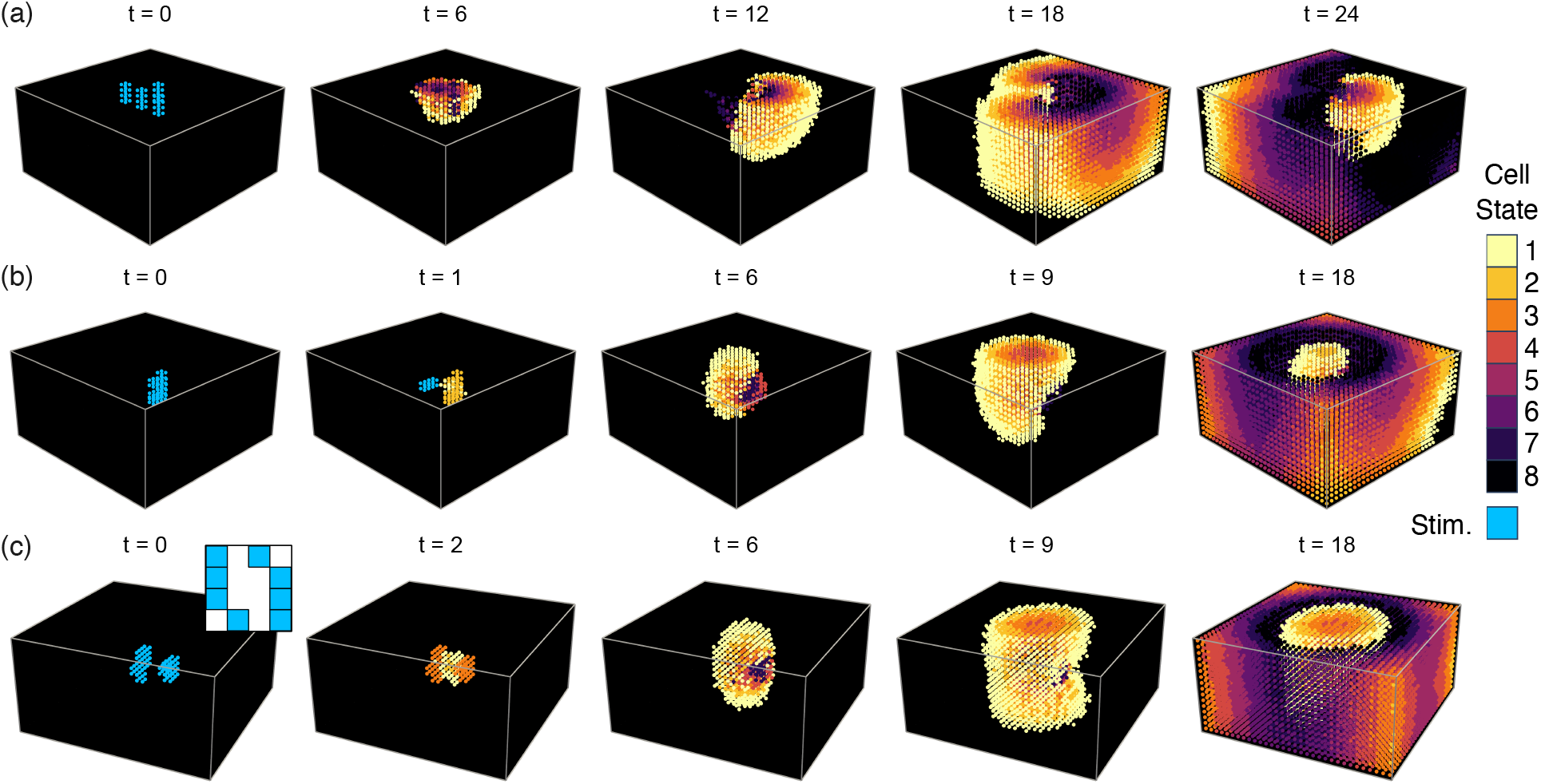
Reentry in the three-dimensional cube cellular-automaton geometry. (a) Four-site spatial subthreshold stimulation pattern from Fig. 5a, applied to the upper surface of the cube and extending to a depth of four cells. Cell color denotes the cellular state: active (1–3) or refractory (4–8). Blue indicates externally stimulated cells. (b,c) Stimulation protocols learned by the RL agent when stimulation is restricted to a central region of the cube. (b) Sequential subthreshold stimulation initiates unidirectional propagation and produces a pair of scroll waves. (c) Spatial subthreshold stimulation consisting of a single ringlike stimulus in an *x*–*y* plane near the center of the cube. The inset shows the stimulus projected onto the *x*–*y* plane. The stimulus generates inward propagation toward the center of the ring, followed by propagation above and below the stimulated plane, resulting in two pairs of scroll waves.

Training the RL agent directly on the threedimensional geometry revealed strategies for initiating reentry from the center of the cube. A frequent solution consisted of two sequential subthreshold pulses (Fig. 7b), resulting in unidirectional propagation and the formation of a pair of scroll waves. This solution can therefore be understood as a three-dimensional realization of the sequential subthreshold mechanism identified in two dimensions (Fig. 5b).

The highest-reward solution used the spatial subthreshold mechanism, but in a form adapted to the three-dimensional geometry (Fig. 7c). A single stimulus pulse, arranged as a ring-like pattern in an *x*–*y* plane near the center of the cube, generated propagation inward toward the center of the ring. The wave then propagated above and below the stimulated plane, before curling around the refractory region created by the initial stimulus. Removal of any one stimulation site resulted in propagation block, except for one site, whose removal produced unidirectional upward propagation only. This solution generated greater sustained activity than the two-pulse protocol and therefore received higher reward.

## 6 Experimental Observations

We next investigated whether the spatial subthreshold mechanism identified by the RL agent could be observed in cardiac tissue. We focused on the four-site stimulus pattern shown in Figs. 5a and 7a. A defining feature of this mechanism is that wave initiation arises from the combined effect of several individually subthreshold stimulus sites: each site is insufficient to initiate propagation on its own, but together the sites generate a propagating wave. To test this principle experimentally, we used optogenetic stimulation in cardiac preparations expressing channelrhodopsin-2 (ChR2), which locally depolarizes tissue in response to patterned blue light (see Appendix B).

In each experiment, the light intensity and spot spacing were adjusted so that removing any one spot abolished propagation, whereas the complete four-site pattern produced excitation that first appeared between two or more illuminated regions. The spacing was selected using the same functional criterion as in the cellular automaton: the stimulus sites were positioned so that their combined activity generated excitation in the intervening region, while each site remained individually subthreshold. The model and experimental patterns were therefore comparable in terms of the spatial interaction required for multisite excitation.

We first tested the mechanism in two-dimensional cardiac monolayers. These preparations consist of thin cultured sheets of interconnected cardiac cells that support undamped wave propagation in two dimensions and can be virally transduced to express ChR2 [52]. Four-site patterns were projected onto the monolayers, with the relative spacing, orientation, and intensity of the spots adjusted to produce near-threshold excitation. To generate repeatable experiments in uniformly repolarized tissue, the monolayer was illuminated with a long depolarizing pulse, followed by the patterned stimulus at a fixed interval (see Appendix B.1). Under these conditions, waves initiated between illuminated regions, consistent with excitation arising from the combined depolarizing effect of multiple subthreshold sites rather than from any individual site alone. Depending on the spacing and orientation of the pattern, the resulting waves either failed to propagate, formed target waves, or initially propagated unidirectionally before resolving into either target waves or spiral waves (Fig. 8a–h).

**Fig 8.**
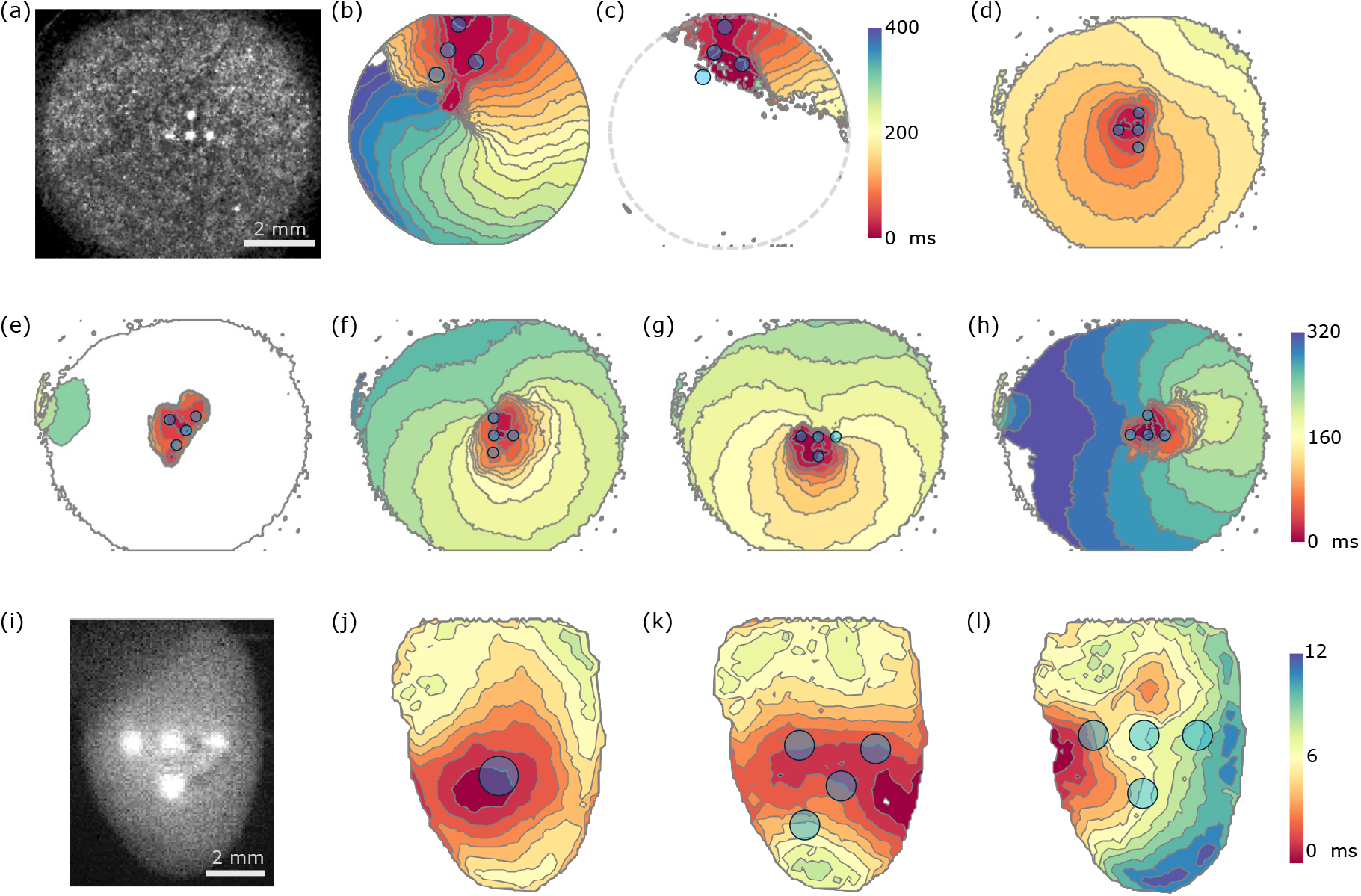
Experimental results in two-dimensional cardiac monolayers and intact hearts. (a) Optogenetically sensitized cardiac monolayer exposed to patterned blue-light stimulation, with the stem of the four-site T-shaped pattern oriented upward (see Appendix B.1). (b,c) When positioned near the edge of the monolayer, the pattern generates a unidirectional wave that propagates clockwise and completes either a full circuit (3 of 5 trials) or a partial circuit (2 of 5 trials). (d–h) Rotation of the four-site pattern produces different outcomes, including (d) target waves, (e) propagation block, and (f–h) initially unidirectional propagation. In each unidirectional case, the wave initially propagates within the 180^°^ sector centered on the direction from the central stimulus site toward the base of the T-shaped pattern. (i–l) Patterned optical stimulation of optogenetically sensitized intact hearts (see Appendix B.2). A single large stimulus generates a target wave (j), whereas four-site patterns can generate waves that originate between or adjacent to multiple stimulation sites (k). In some cases, the resulting wave initially propagates preferentially in one direction before colliding with other wavefronts (l).

The responses were reproducible for a fixed pattern and orientation. Repeated presentation of the same stimulus generated waves that initiated in the same region and propagated in the same direction, indicating that the near-threshold responses were deterministic under the experimental conditions used here. Across 20 measurements in 11 distinct spacings and rotations of the four-site pattern, 30% produced target waves, 20% blocked, and 50% produced initially unidirectional waves. A similar distribution of outcomes was observed in the cellular automaton across 16 rotated and spatially shifted configurations of the action space: 6 produced target waves, 4 blocked, and 6 produced reentry (Fig. 5a).

The direction of unidirectional propagation also showed qualitative agreement with the model. In both simulations and experiments, unidirectional waves initially propagate within the 180^°^ sector centered on the direction from the top-center stimulus site toward the base of the ‘T’. However, rotating the stimulus pattern did not produce an exact corresponding rotation of the wave direction. Instead, different orientations of the same pattern produced different outcomes and propagation directions, but all within the 180^°^ sector defined by the rotated pattern.

As a preliminary extension to three-dimensional tissue, we also applied four-site stimulus patterns to Langendorff-perfused whole hearts from genetically modified mice expressing ChR2 in cardiac muscle (Fig. 8i–l). In these experiments, we stimulated the tissue late in cardiac diastole (Appendix B.2) in order to ensure the tissue was uniformly recovered. The patterns were scaled and the light intensity varied so that excitation was initiated between illuminated regions, as in the monolayer experiments. In some orientations, the early activation pattern was consistent with preferential propagation in one direction before resolving to a target wave; however, these results must be interpreted in light of experimental limitations (see Discussion below).

## 7 Discussion

The RL agent learned several approaches to initiate reentry across different geometries. These approaches share a common feature: each introduces or exploits an asymmetry. The agent introduced asymmetry in stimulus strength (the superthreshold–subthreshold mechanism), asymmetry in timing (sequential subthreshold mechanism), and asymmetry in spatial arrangement (spatial subthreshold mechanism). In more complex domains, the agent also exploited local asymmetries in the geometry, such as boundaries and corners.

The subthreshold stimulation mechanisms identified here reveal a potential physiological pathway by which localized spontaneous cellular activity can initiate reentry. Small regions of spontaneous activity, including activity associated with early afterdepolarizations, can produce localized depolarizations that are individually insufficient to initiate a propagating wave. Our results suggest that the spatial and temporal organization of several such events may nevertheless generate unidirectional propagation and create the conditions for reentry.

Interestingly, the RL agent did not discover the classical mechanism of reentry initiation in which a superthreshold stimulus is applied to the refractory tail of a propagating wave. One possible explanation is that this mechanism is disfavored by the reward function used in this work. Because the reward penalizes the number of stimulated cells, generating a propagating wave with superthreshold stimulation incurs a greater cost than the subthreshold mechanisms identified here. Consequently, the learned policies favor stimulation protocols that initiate reentry with fewer stimulated cells.

Training RL agents typically requires a large number of simulations, which favors computationally tractable models such as cellular automata over detailed biophysical models. This abstraction allowed us to train agents efficiently across multiple geometries and to identify general mechanisms for initiating reentry. At the same time, cellular automata omit biophysical effects that may influence reentry in cardiac tissue. With larger computational budgets, training agents directly on detailed biophysical models could allow RL to discover strategies tied to processes not captured by the cellular automaton, such as action-potential-duration restitution, cellular ionic properties, and alternative mechanisms of intercellular conduction, including ephaptic coupling [53, 54].

We sought experimental confirmation of the mechanisms identified by the agent, first in the 2D monolayer preparation and then in the whole heart. In the monolayer, subthreshold pulses combined to generate a propagating wave, and the resulting dynamics were reproducible (Fig. 9b). This demonstration of repeatable, deterministic behavior supports our strategy of training RL agents to identify reentrant mechanisms in deterministic models of excitable media. The area excited by the spot pattern is also consistent with the model, with total area being less than required by a single superthreshold pulse (Fig. 9c). The experiments reproduced two specific predictions of the simulations: unidirectional waves propagated in a preferred direction relative to the orientation of the stimulation pattern, and the direction of the wavefront showed inherent variability (Fig. 8f–h). The proportion of unidirectional, target, and blocked waves was also consistent between simulation and experiment. These lines of evidence indicate that the monolayer follows mechanisms similar to those discovered by the RL agent.

**Fig 9.**
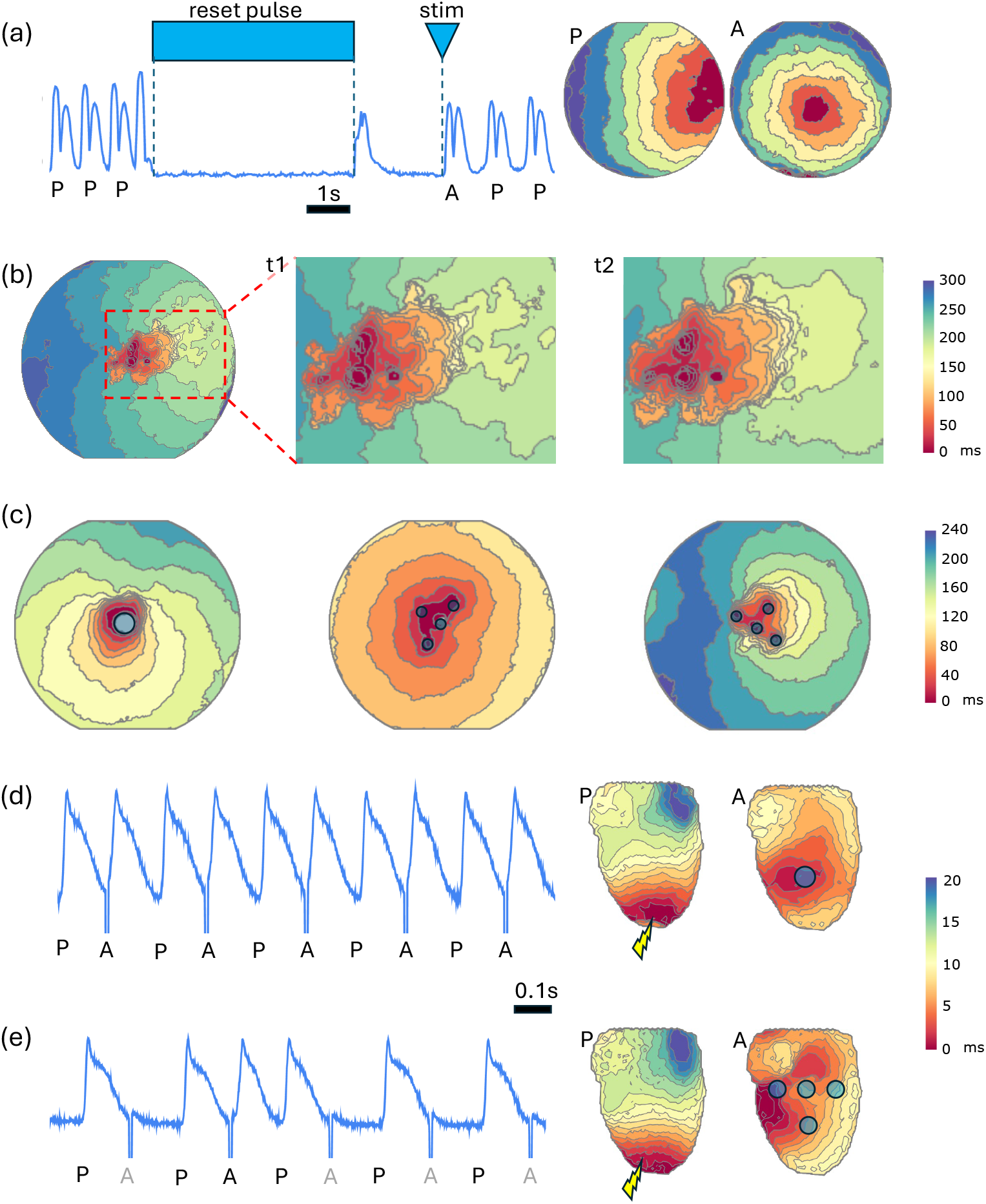
Experimental protocols and supporting data. (a) Monolayer preparations exhibit spontaneous target or reentrant waves. To suppress this intrinsic activity, the entire monolayer is exposed to prolonged depolarizing illumination (the ‘reset pulse’), followed after a fixed interval by a brief patterned stimulus (‘stim’). The pulse durations and timing are chosen so that the patterned stimulus is delivered shortly before spontaneous activity resumes. Here, ‘P’ denotes intrinsic pacemaker activation and ‘A’ denotes activation by patterned light. In this example, the intrinsic pacemaker is located on the right, and a single circular spot projected at the center generates a target wave. (b) Two trials (t1 and t2) using the same stimulation pattern and protocol produce similar activation maps. (c) Comparison of waves generated by a single large stimulus (left; one 0.9-mm-diameter spot) and four smaller stimuli (middle and right; four 0.35-mm-diameter spots). Target waves propagate in all directions, producing isochrones that form closed rings (left and middle), whereas a unidirectional wave initially propagates preferentially in one direction, producing isochrones that form broken rings (right). (d,e) Whole-heart stimulation protocol. The heart is electrically paced at the apex with a superthreshold pulse (‘P’ in the trace), followed after a fixed interval by patterned optical stimulation (‘A’). (d) A superthreshold optical stimulus consistently generates a target wave centered on the stimulated region. (e) Patterns of subthreshold spots generate wavefronts intermittently. For some orientations of the four-site pattern, excitation begins between or adjacent to multiple stimulation sites and initially propagates preferentially in one direction.

The whole-heart experiments provide a preliminary extension of this result to intact threedimensional tissue. Four-site patterns produced excitation waves that originated between illumination spots and, in some cases, propagated preferentially in one direction at onset before resolving to target waves (Fig. 8k–l). These observations suggest that multisite subthreshold stimulation can shape early propagation in the intact heart. However, validating the specific patterns identified by the RL agent is more challenging in the whole-heart preparation than in the monolayer.

First, although all sites in a pattern are illuminated simultaneously for 3 ms, ChR2 channels remain open for 15–30 ms after light offset, with longer closing times at lower light intensities and depolarized voltage [55]. When coupled with heterogeneity in ChR2 expression [56], this may result in sites depolarizing and reaching threshold at different rates. These differences will have greater impact on wave position in whole-heart preparations as they have significantly higher conduction velocities. Second, the effective spacing between illumination sites in intact heart tissue is determined not only by their distance on the epicardial surface, but also by fiber anisotropy [57], transmural anatomy, depth-dependent light penetration, and local recovery state [58, 59]. Consequently, rotating the same projected pattern changes its functional geometry relative to the tissue: a pattern that is subthreshold at each individual site in one orientation may behave as effectively superthreshold in another, making pattern rotation an unsuitable controlled test of directional dependence. In addition, we observed that near-threshold stimuli generated waves intermittently. While sporadic activation is expected for stimuli near threshold [60], it resulted in variations in action potential duration (Fig. 9d,e) which impacted the recovery state of the tissue and also could lead to recovery-dependent changes in tissue space constant [58]. Future experiments will require training RL environments on realistic ventricular geometries incorporating anisotropic conduction, optical penetration, and heterogeneous excitability.

Reinforcement learning has also begun to be explored as a strategy for terminating established reentrant activity. Previous work has used reinforcement learning to steer an idealized model of a spiral-wave core toward a boundary, where it was eliminated [61], and to design ablation strategies for cardiac-tissue models and patient-specific image-based atrial models [62, 63]. Building on these studies, the framework developed here could be adapted to optimize the timing, location, and geometry of subthreshold pulses for terminating reentry. Such approaches could complement established low-energy antifibrillation pacing strategies and have relevance to cardioversion, antitachycardia pacing, and targeted ablation [64, 65].

The initiation of reentry is a fundamental problem in the physics of excitable media and a central question in cardiac electrophysiology, with proposed mechanisms spanning cellular, tissue-level, and geometrical effects. Our results demonstrate that RL can discover distinct initiation mechanisms across different geometries, including mechanisms based on stimulus strength, timing, spatial arrangement, and structural asymmetry. As such, RL provides a promising framework for generating testable hypotheses about the initiation of reentry in cardiac tissue and other excitable systems, with the potential to guide new experimental designs and clinical investigations. Just as RL has discovered strategies in Go and chess that surprised even expert human players [39], it may uncover similarly unexpected mechanisms of reentry in excitable systems.

## Acknowledgements

T.M.B. acknowledges support from a Fonds de recherche du Québec – Nature et technologies (FRQNT) postdoctoral fellowship, during which part of this work was completed. G.P. acknowledges support from the Mackey–Glass Research Bursary in Physiology. G.B. acknowledges support from the Heart and Stroke Foundation of Canada (G-18-0022123) and the Natural Sciences and Engineering Research Council of Canada (NSERC; RGPIN-2018-05346). This research was supported in part by grant NSF PHY-2309135 to the Kavli Institute for Theoretical Physics (KITP) and the Gordon and Betty Moore Foundation Grant No. 2019.02. Computational resources were provided in part by Calcul Québec and the Digital Research Alliance of Canada. We thank Flavio Fenton, Abouzar Kaboudian and Leon Glass for helpful discussions, and Abouzar Kaboudian for testing stimulation patterns in abubu.js.

We thank Lorenzo Rezza for his contribution to the development of the optical mapping software, Natasha Giacomelli for her contribution to the animal procedures, Leslie M. Loew for providing the ElectroFluor 730P voltage-sensitive dye, and Marina Campione for providing the ChR2 mouse line.

## Data availability

Code for the cellular automaton simulations, RL environments, training scripts, and figure generation is available from Ref. [66]. The version corresponding to this article is archived on Zenodo at Ref. [67]. The experimental data are available from the authors upon reasonable request.

## Appendices

### A Reaction–diffusion model simulations

We tested the learned stimulation mechanisms in three reaction–diffusion models of excitable media: the FitzHugh–Nagumo, Fenton–Karma [49], and O’Hara–Rudy models [50]. All simulations were performed on one-dimensional cables using Myokit [48]. The Laplacian was discretized on a regular grid. Since the spatial scale is arbitrary in these simulations, changing the diffusive coupling strength is equivalent to rescaling the cable length.

#### A.1 FitzHugh–Nagumo model

We simulated the FitzHugh–Nagumo reaction– diffusion model,

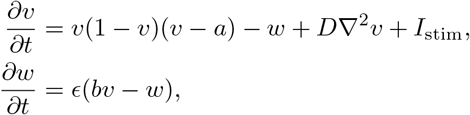

where *v* is the fast excitation variable, *w* is the slow recovery variable, *D* is the diffusion coefficient, and *I*_stim_ is an externally applied stimulus. The model is nondimensional and phenomenological. Accordingly, space, time, the excitation and recovery variables, and stimulus amplitude are expressed in arbitrary units. We used parameters

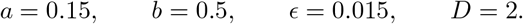

Simulations were performed on a cable of 128 grid points and integrated with time step Δ*t* = 0.01. Stimuli were applied by setting *I*_stim_ = 0.5 within the specified stimulation region for a duration of one time unit, with *I*_stim_ = 0 elsewhere.

#### A.2 Fenton–Karma model

Simulations were performed on a cable of length 1024 grid cells using parameter set 4 described by Fenton et al. [68] and diffusive coupling conductance 20. The model was prepaced for 1000 beats at a basic cycle length of 1000 ms before applying the stimulation protocol. Simulations were integrated with time step Δ*t* = 0.01 ms. Stimuli were applied by setting *I*_stim_ = 50 mV*/*ms within the specified stimulation region for a duration of 1 ms.

#### A.3 O’Hara–Rudy model

Simulations were performed on a cable of length 512 grid cells using the default model parameters and diffusive coupling conductance 2. The model was prepaced for 1000 beats at a basic cycle length of 1000 ms before applying the stimulation protocol. Because of the rapid upstroke, simulations were integrated with time step Δ*t* = 10^−4^ ms. Stimuli were applied by setting *I*_stim_ = 40 *µ*A*/µ*F within the specified stimulation region for a duration of 1 ms.

### B Experimental setup

#### Ethics statement

The use of postnatal mouse cardiac tissue for the preparation of ventricular monolayers was conducted in compliance with Canadian Council on Animal Care guidelines and institutional regulations and was reviewed and approved by the McGill University Animal Care Committee under protocol 2018–8044 (SOP 301–01). All procedures involving the whole-heart experiments were performed in accordance with Directive 2010/63/EU of the European Parliament on the protection of animals used for scientific purposes and were approved by the Italian Ministry of Health under protocol 944/2018-PR.

#### B.1 Monolayer experiments

We prepared 8-mm-diameter monolayers of cardiac cells from neonatal mouse ventricles and genetically modified them to express the light-sensitive ion channel channelrhodopsin-2 (ChR2), which enables optical control of cardiac excitation [69]. Experiments were performed in a stage-top CO_2_ incubator at 33–35^°^C using optical stimulation and dye-free motion detection, as first described in Ref. [52]. Images were acquired using a machine-vision camera (Basler acA1920-155um) at 50 frames per second. Multisite stimulation patterns were rescaled and the diameter of each spot was adjusted during experiments so that the following conditions were satisfied: (i) removal of any one spot resulted in no propagating wave, (ii) increasing the spot diameter or adding an additional spot resulted in a target wave, and (iii) wave fronts were initiated between illumination spots.

To suppress spontaneous activity, the entire tissue was first depolarized using uniform blue-light illumination for several seconds. Following termination of this reset pulse, the monolayer remained quiescent for a short period. The duration of this quiescent period was determined experimentally for each monolayer, allowing subsequent test stimuli (e.g., the four-site pattern) to be delivered in uniformly recovered tissue.

We also experimentally verified two additional features corresponding to conditions examined in the simulations. First, preparations were checked for stability under this protocol by confirming that propagation patterns elicited by the same four-site stimulus were reproducible across repeated trials (Fig. 9b). Second, we confirmed that the total illuminated area of the four-site pattern was smaller than that of a single circular superthreshold stimulus (Fig. 9c); specifically, the total illuminated area of the four-spot pattern was approximately half that of a single circular stimulus just above the minimum diameter required for propagation.

For the experiments shown in Fig. 8, the reset pulse lasted 8 s, and an 80-ms stimulus was delivered 1.5 s after the end of the reset pulse. For the four-site patterns shown in Figs. 8d–h and 9, the inter-spot distance and spot diameter were 0.9 mm and 0.35 mm, respectively. For Figs. 8b,c, the corresponding values were 1.1 mm and 0.5 mm.

#### B.2 Whole-heart experiments

We performed experiments on three mice with genetically modified hearts expressing channelrhodopsin-2 (ChR2; see Ref. [70]). Mice were anesthetized with inhaled isoflurane, after which the hearts were rapidly excised, immersed in Krebs–Henseleit (KH) solution, and cannulated through the aorta. Contraction was inhibited by adding 10 *µ*M (±)-blebbistatin to the perfusion solution. The cannulated hearts were then transferred to the recording chamber and retrogradely perfused using a Langendorff apparatus with KH solution at a constant flow rate of 2.5 ml/min. After several minutes of perfusion, 1 ml of KH solution containing the voltage-sensitive dye ElectroFluor 730p at 8 *µ*g/ml was injected as a bolus through the aorta [71].

For imaging, we used an optical-mapping system previously described in Ref. [72]. Briefly, the whole heart was illuminated using a light-emitting diode operating at a wavelength centered at 730 nm. Fluorescence emission was collected through a dichroic beam splitter and a 792/64-nm bandpass filter and recorded at a spatial resolution of 128 × 128 pixels and a frame rate of 1000 frames per second. The size and intensity of the optical stimulation patterns were adjusted using the same procedure as in the monolayer experiments.

To deliver optical stimulation at a consistent phase of the cardiac cycle, we used a protocol similar to that of Ref. [58]. The heart was first paced at the apex using a superthreshold electrical stimulus, followed after a fixed interval by a patterned optical stimulus applied to the ventricles. The interval between the two stimuli was selected such that electrically and optically induced waves occurred in alternation (Fig. 9d,e). The interspot distance used in the whole-heart experiments was 1.3 mm.

