## Supplementary Information for "Reinforcement learning discovers new mechanisms of reentry in excitable media"

### 1 Simulation of reaction–diffusion models

#### 1.1 FitzHugh–Nagumo model simulations

We simulated the FitzHugh–Nagumo reaction–diffusion model,

$$\frac{\partial v}{\partial t} = v(1-v)(v-a) - w + D\nabla^2 v + I_{\text{stim}}, \tag{1}$$

$$\frac{\partial w}{\partial t} = \epsilon(bv - w), \tag{2}$$

where $v$ is the fast excitation variable, $w$ is the slow recovery variable, $D$ is the diffusion coefficient, and $I_{\text{stim}}$ is an externally applied stimulus current. We used parameter values

$$a = 0.15, \qquad b = 0.5, \qquad \epsilon = 0.015, \qquad D = 2.$$

The parameter $\epsilon$ controls the separation of time scales between the fast excitation and slow recovery variables. The diffusion coefficient $D$ sets the spatial scale of propagation; since the spatial scale is arbitrary in these simulations, changing $D$ is equivalent to rescaling the cable length.

Simulations were performed on a one-dimensional cable represented by 128 grid cells using Myokit [1]. The Laplacian was discretized on this one-dimensional grid, and the equations were integrated with a time step of $\Delta t = 0.01$. Stimuli were applied by setting $I_{\text{stim}} = 0.5$ within the specified stimulation region for a duration of one time unit, with $I_{\text{stim}} = 0$ elsewhere.

#### 1.2 Fenton–Karma model simulations

We also tested the stimulation mechanisms in the Fenton–Karma model [2], using the implementation provided in Myokit [1]. Simulations were performed on a one-dimensional cable represented by 1024 grid cells. As in the FitzHugh–Nagumo simulations, the spatial scale is arbitrary; changing the diffusive coupling strength is therefore equivalent to rescaling the cable length.

We used Fenton–Karma parameter set 4, with a diffusive coupling conductance of 20. The model was prepaced at a basic cycle length of 1000 ms before applying the stimulation protocol. Simulations were integrated with time step figure$\Delta t = 0.01$ ms. Stimuli were applied by setting $I_{\text{stim}} = 50$ mV/ms within the specified stimulation region for a duration of 1 ms.

#### 1.3 O'Hara–Rudy model simulations

We further tested the stimulation mechanisms in the O'Hara–Rudy dynamic human ventricular model [3], using the implementation provided in Myokit [1]. Simulations were performed on a one-dimensional cable represented by 512 grid cells. As in the other reaction–diffusion simulations, the spatial scale is arbitrary; changing the diffusive coupling strength is therefore equivalent to rescaling the cable length.

We used the default O'Hara–Rudy model parameters, with a diffusive coupling conductance of 2. Because of the rapid upstroke of the O'Hara–Rudy model, simulations were integrated using a time step of $\Delta t = 10^{-4}$ ms.

Before applying the stimulation protocol, the model was prepaced for 1000 beats at a basic cycle length of 1000 ms. Stimuli were applied by setting $I_{\text{stim}} = 40$ $\mu$A/$\mu$F within the specified stimulation region for a duration of 1 ms.

### 2 Supplementary Figures

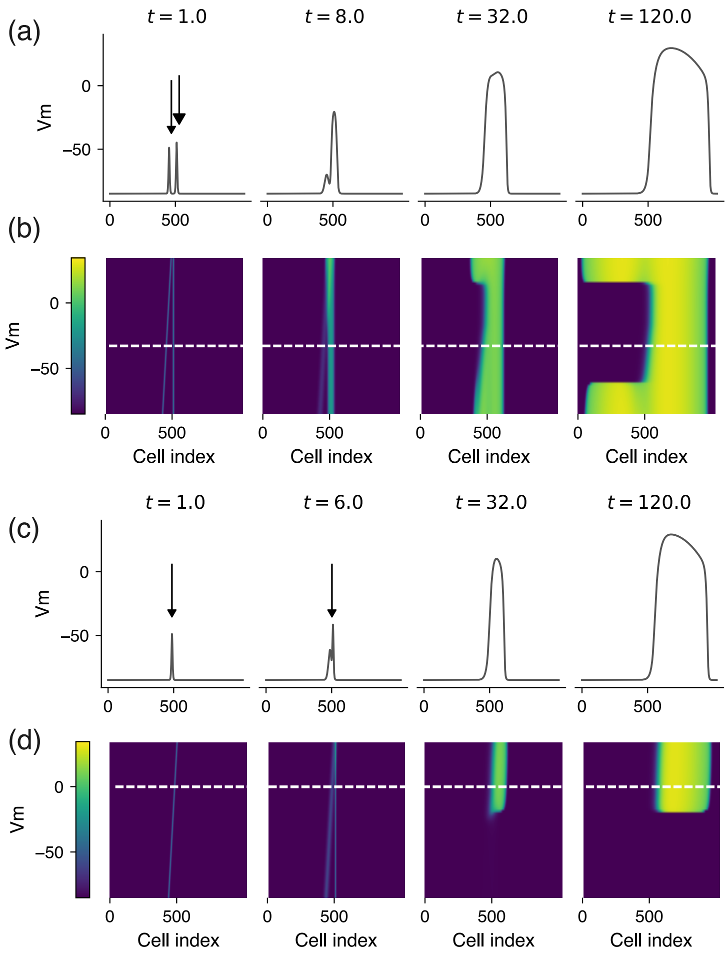


**Fig. 1** Initiation of unidirectional propagation in a cable represented by 1024 grid cells described by the Fenton–Karma model. (a) Voltage profiles as a function of space at selected times. At $t = 0$, stimuli of duration 1 ms are applied to two spatial regions separated by 50 cells: one spanning 7 cells (subthreshold, small arrowhead) and the other spanning 8 cells (superthreshold, large arrowhead). (b) Voltage profiles from 32 vertically stacked cables (hence propagation only in the horizontal direction), illustrating the effect of varying the separation between the subthreshold and superthreshold stimuli. The white dashed line indicates the cable shown in (a). (c) Unidirectional propagation induced by two sequential subthreshold stimuli. The stimuli are applied 5 ms apart to regions spanning 7 cells each, separated by 16 cells. (d) Voltage profiles from 32 stacked cables showing the effect of varying the spatial separation between the two subthreshold stimuli.

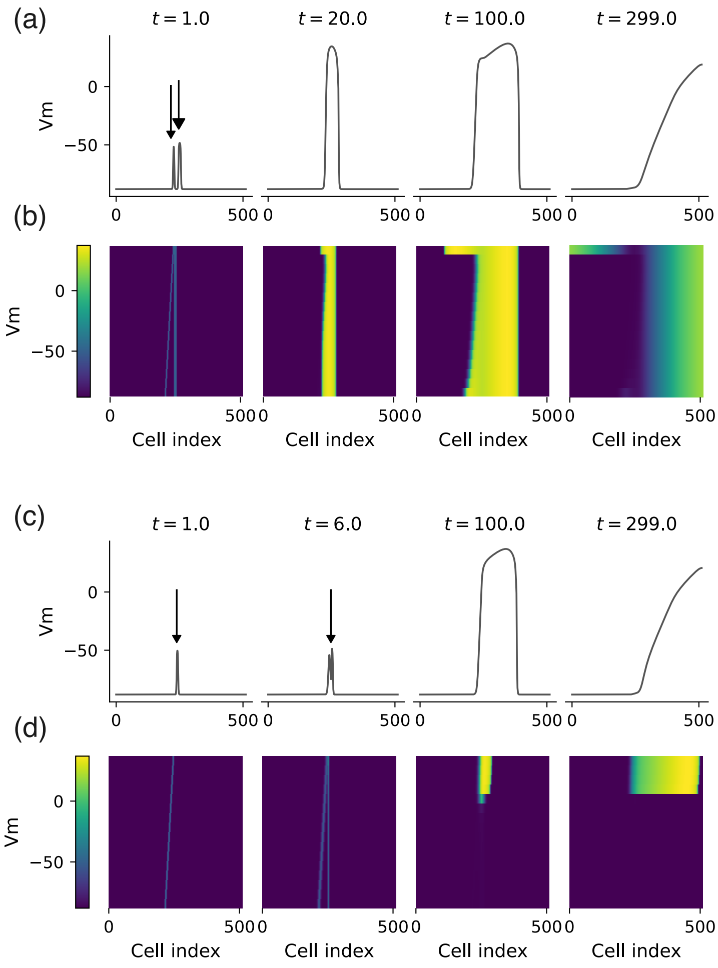


**Fig. 2** Initiation of unidirectional propagation in a cable represented by 512 grid cells described by the O'Hara–Rudy model. (a) Voltage profiles as a function of space at selected times. At $t = 0$, stimuli of duration 1 ms are applied to two spatial regions separated by 16 cells: one spanning 5 cells (subthreshold, small arrowhead) and the other spanning 10 cells (superthreshold, large arrowhead). (b) Voltage profiles from 32 vertically stacked cables (hence propagation only in the horizontal direction), illustrating the effect of varying the separation between the subthreshold and superthreshold stimuli. The white dashed line indicates the cable shown in (a). (c) Unidirectional propagation induced by two sequential subthreshold stimuli. The stimuli are applied 5 ms apart to regions spanning 6 cells each, separated by 5 cells. (d) Voltage profiles from 32 stacked cables showing the effect of varying the spatial separation between the two subthreshold stimuli.
